# Prediction of celiac disease associated epitopes and motifs in a protein

**DOI:** 10.1101/2022.07.26.501507

**Authors:** Ritu Tomer, Sumeet Patiyal, Anjali Dhall, Gajendra P. S. Raghava

**Author notes:** **Corresponding Author** Prof. Gajendra P. S. Raghava, Head and Professor, Department of Computational Biology, Indraprastha Institute of Information Technology, Delhi, Okhla Industrial Estate, Phase III, (Near Govind Puri Metro Station), New Delhi, India – 110020 Office: A-302 (R&D Block), Website: http://webs.iiitd.edu.in/raghava/. **Mailing Address of Authors** Ritu Tomer (RT) Sumeet Patiyal (SP) Anjali Dhall (AD) Gajendra P. S. Raghava (GPSR). Equal Contribution (Ritu Tomer, Sumeet Patiyal, Anjali Dhall).

## Abstract

Celiac disease (CD) is an autoimmune gastrointestinal disorder which causes immune-mediated enteropathy against gluten. The gluten immunogenic peptides have the potential to trigger immune responses which leads to damage the small intestine. HLA-DQ2 and HLA-DQ8 are major alleles that bind to epitope/antigenic region of gluten and induce celiac disease. There is a need to identify CD associated epitopes in protein-based foods and therapeutics. In addition, prediction of CD associated epitope/peptide is also required for developing antigen-based immunotherapy against celiac disease. In this study, computational tools have been developed to predict CD associated epitopes and motifs. Dataset used in this study for training, testing and evaluation contain experimentally validated CD associated and non-CD associate peptides. Our analysis support existing hypothesis that proline (P) and glutamine (Q) are highly abundant in CD associated peptides. A model based on density of P&Q in peptides has been developed for predicting CD associated which achieve maximum AUROC 0.98. We discovered CD associated motifs (e.g., QPF, QPQ, PYP) which occurs specifically in CD associated peptides. We also developed machine learning based models using peptide composition and achieved maximum AUROC 0.99. Finally, we developed ensemble method that combines motif-based approach and machine learning based models. The ensemble model-predict CD associated motifs with 100% accuracy on an independent dataset, not used for training. Finally, the best models and motifs has been integrated in a web server and standalone software package “CDpred”. We hope this server anticipate the scientific community for the prediction, designing and scanning of CD associated peptides as well as CD associated motifs in a protein/peptide sequence (https://webs.iiitd.edu.in/raghava/cdpred/).

**Key Points:**

- Celiac disease is one of the prominent autoimmune diseases
- Gluten immunogenic peptides are responsible for celiac disease
- Mapping of celiac disease associated epitopes and motifs on a proteins
- Identification of proline and glutamine rich regions
- A web server and software package for predicting CD associate peptides

**Author’s Biography:**

1. Ritu Tomer is currently working as Ph.D. in Computational Biology from Department of Computational Biology, Indraprastha Institute of Information Technology, New Delhi, India.
2. Sumeet Patiyal is currently working as Ph.D. in Computational biology from Department of Computational Biology, Indraprastha Institute of Information Technology, New Delhi, India.
3. Anjali Dhall is currently working as Ph.D. in Computational Biology from Department of Computational Biology, Indraprastha Institute of Information Technology, New Delhi, India.
4. Gajendra P. S. Raghava is currently working as Professor and Head of Department of Computational Biology, Indraprastha Institute of Information Technology, New Delhi, India.

## Introduction

Celiac disease (CD) is an auto-immunological disorder which mainly affects the small intestine of the infected person [1]. CD is a life-long disorder occurred due to the gluten associated foods which is found in various foods such as wheat, barley, spelt, kamut, and rye [2]. The prevalence rate of CD is around 1.4% worldwide and it may vary with genetic and environmental factors. The occurrence of disease is significantly higher in children in comparison to adults [3]. The presence of human leukocyte antigens (HLAs) play a crucial role in the induction and regulation of immunological responses [4]. The gluten immunogenic peptides bind with specific MHC class II binders i.e., HLA-DQ2/HLA-DQ8 in lamina propria region and activate both innate and adaptive immune system [5]. Studies shows that HLA-DQ2 found in almost 94.5% of CD cases while HLA-DQ8 present in 2.7% of the cases [6]. These binders are also associated with other autoimmunological disorders such as HLA-DQ8 associated with Type I diabetes [7] while HLA-DQ6 and DQ8 are associated with Multiple sclerosis [8].

The entry of gluten inside the lamina propria region of small intestine follows majorly two pathways, transcellular pathway and paracellular pathway [9], depicted in Figure 1. In transcellular pathway, the entry of gluten is associated with the binding of secretory IgA (sIgA) in the apical region of intestine [10]. While in the paracellular pathway, the entry of gluten is associated with the binding of chemokine receptor 3 (CXCR3) present at enterocyte with the release of zonulin protein [11,12]. After entering inside the lamina propria region, a series of events occurs to an inflammatory cascade which leads to damaging the intestinal villi by the excessive release of antibodies (anti-tissue transglutaminase (tTG2), anti-IgA antibodies and anti-endomysial antibodies) and cytokine [13].

**Figure 1:**
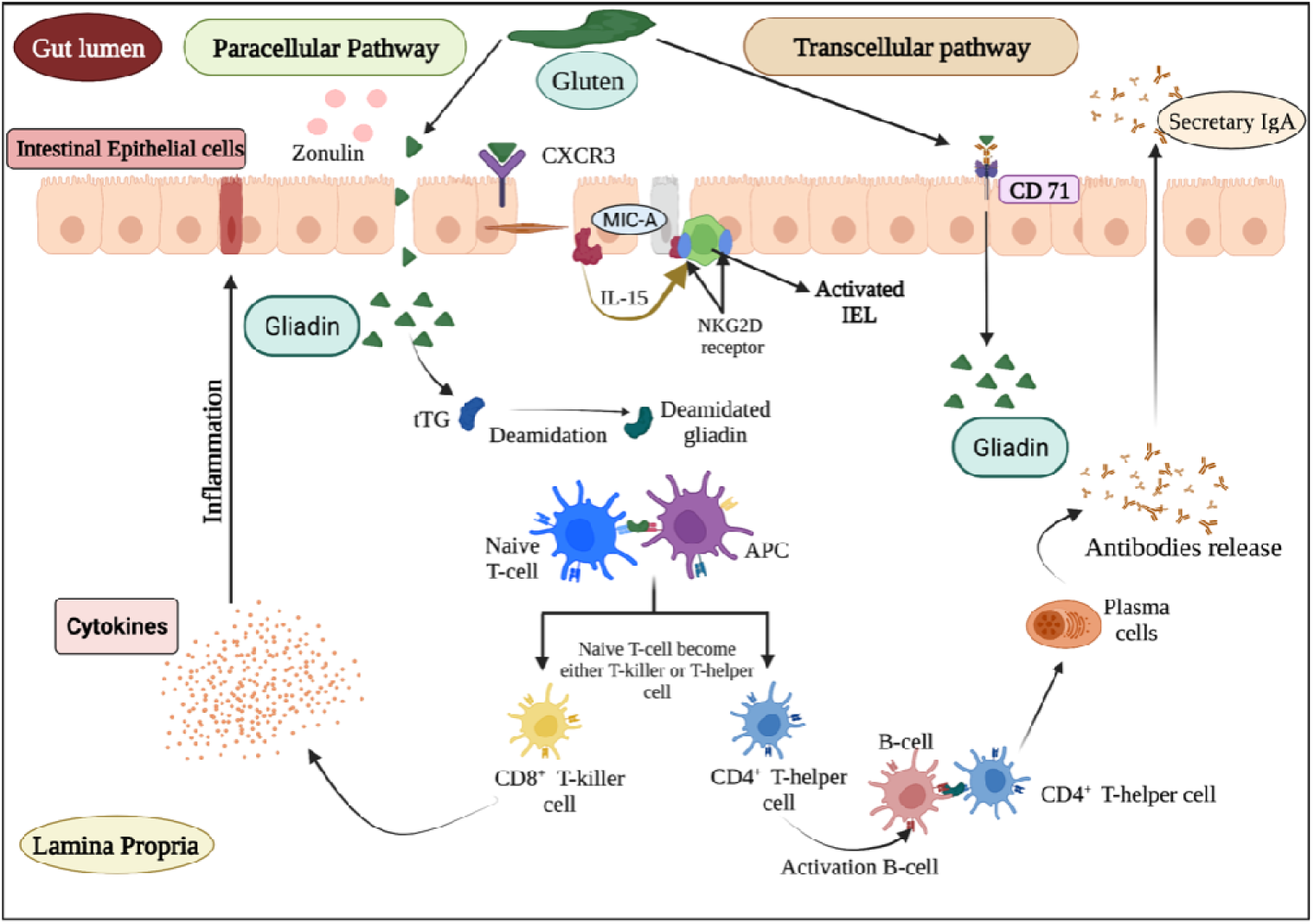
Schematic representation of celiac disease pathogenesis and immune response.

Due to auto-inflammatory immune responses several gastrointestinal disorders like malabsorption, vomiting, bloating, diarrhoea, abdominal pain and distension occurred [14]. Recently, a number of biological and genetic tests (such as detection of antibodies, intestinal tissue biopsy, HLA-typing and gluten challenge test) are available for the disease detection [1]. It has been found in many studies that α-gliadin 33-mer peptide having the property of resistant to gastrointestinal cleavage and makes it highly immunogenic peptide [15–18]. Despite tremendous understanding of CD, effective treatment for the disease is life-long gluten free diet. In order to manage severity of CD effectively, it is important to identify CD associated epitopes or immunogenic peptides responsible for CD. Identification of CD associated epitopes/peptides is not only important for identifying CD free food/therapeutic proteins, it is also important for designing antigen-based immunotherapy against CD.

In the pilot study, we have developed a computational approach for the prediction of CD associated peptides. We have extracted the experimentally validated gluten immunogenic peptides responsible for CD from the IEDB database. In order to create negative dataset, we have collected CD non-causing peptides and random peptides from IEDB and Swiss-Prot, respectively. We have identified highly conserved regions of disease-causing peptides using motif-based search. In addition, we have developed prediction models using composition-based features and machine learning algorithms. In order to facilitate the community, we have provided the webserver and standalone package for the prediction and scanning of CD causing protein/peptides using sequence information.

## Material & Methods

The complete architecture of our study is illustrated in Figure 2. The detail of each step is described below.

**Figure 2:**
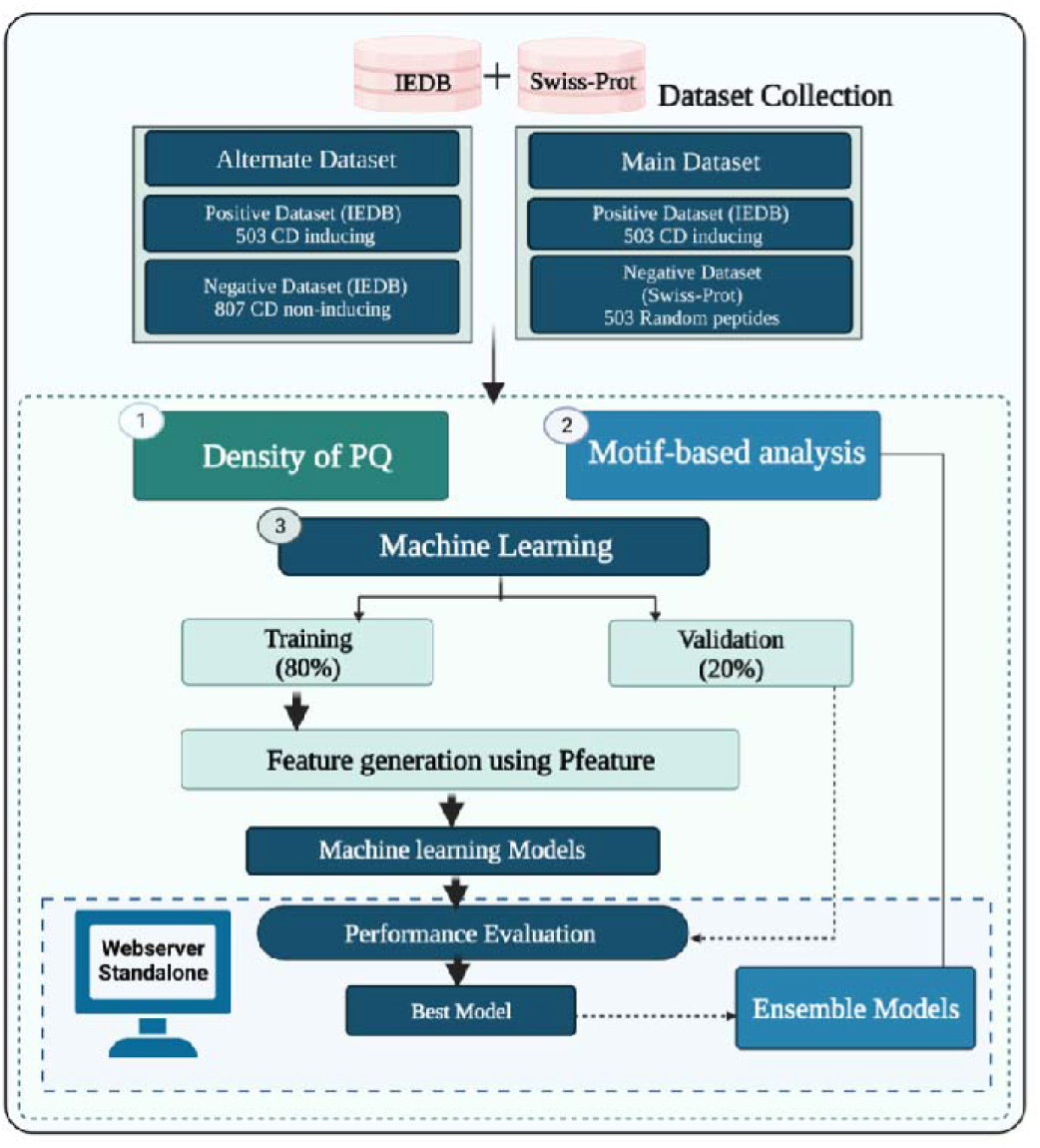
Overall architecture of the study.

### Dataset collection and pre-processing

In this study, we have collected experimentally validated peptides from the immune epitope database (IEDB) [19]. At first, we extracted a total of 521 unique CD causing/associated (gluten immunogenic peptides) from IEDB as a positive dataset. Further, we have selected unique peptides with a length of 9-20 amino-acid residues and got 503 CD associated peptides. Secondly, we extracted experimentally validated CD non-associated peptides from IEDB and random peptides from Swiss-Prot database [20]. The main dataset incorporates 503 CD associated called positive peptides and 503 random peptides called negative peptides. The alternate dataset consists of 503 CD associated and 807 non-associated peptides (which can cause autoimmune disorders other than celiac disease). Finally, we obtained two datasets, i.e., the main dataset comprises an equal number of positive and negative peptides and alternate dataset 503 positive and 807 negative peptides.

### Sequence Logo

In order to understand the preference of amino-acid residues at a specific position, we have generated a one sample logo using WebLogo software [21]. This tool needs a fixed length input sequence vector. Since, the minimum length of peptides in our datasets is 9 residues, so we have extracted 9-mers from N-terminal and 9-mers from C-terminal from each peptide. After that, we re-join both the regions in order to create a fixed length vector of 18 amino acids. The sequences of 18-residues were generated for all the peptides of both positive and negative datasets and used for the creation of one sample logo plots.

### Amino-acid Composition

We have used Pfeature software [22] for the computation of composition-based features. In this current study, we have computed amino acid composition based (AAC) features. In the case of AAC, the composition of each residue is computed in the peptide sequences and a vector od 20 length is generated using the equation 1.

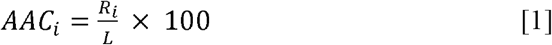

Where, *AAC*_*i*_ is amino-acid composition of residue type *i, R*_*i*_ is the number of residues in *i*, and *L* is the length of peptide sequence.

### Machine Learning Models

We have employed a number of machine learning algorithms for the classification of CD-causing peptides. Currently, we have used scikit-learn [23] python library for the implementation of several classifiers including Decision Tree (DT), Random Forest (RF), XGBoost (XGB), Gaussian Naïve Bayes (GNB) Logistic Regression (LR), ExtraTree classifier (ET), and k-nearest neighbors (KNN).

### Five-fold Cross Validation

In order to avoid overfitting we have train, test and validate the machine learning models by employing five-fold cross validation technique as implemented in previous studied [4,24–27]. At first, the complete dataset was divided into 80:20 ratio, where 80% dataset used for the training and 20% used for the external validation [28–30]. The five-fold cross-validation process is implemented on the 80% training dataset. In this process, the entire training dataset was divided into five equal sets, where each set is used for training and validation purpose. At first, four sets were used for training and fifth set was used for the testing, similarly the process is repeated five times so that each set can be used as testing dataset. Finally, we calculated the average performance of five sets which resulted after five iterations.

### Model Evaluation

In this study, we have used standard parameters for the evaluation of prediction models. Here, we have calculated both threshold dependent as well as independent parameters as previously used in various studies [25]. In the case of threshold-dependent parameters we have computed, sensitivity (Sens), specificity (Spec), accuracy (Acc) and Matthews correlation coefficient (MCC) using the following equations (1-4). In addition, we have measured the performance of models with a well-established and threshold-independent parameter Area Under the Receiver Operating Characteristic (AUROC) curve.

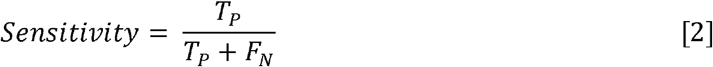

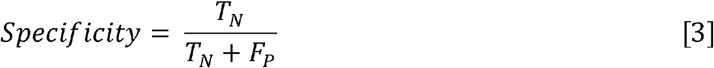

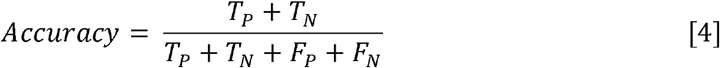

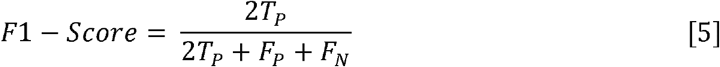

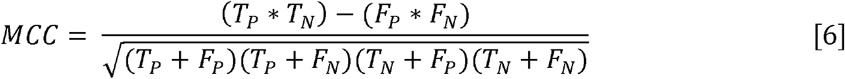

Where, T_P_, T_N_, F_P_ and F_N_ stand for true positive, true negative, false positive and false negative, respectively.

### Ensemble method

The ensemble method is a hybrid approach in which both motifs based, and machine learning methods combined to achieve better performance. In this method, first motif-based approach is used to identify the disease-causing peptides and then we use machine learning methods to predict those peptides which are not covered by the motif-based approach. Finally, we generate an ensemble method which is a combination of both motif-based approach and machine learning method.

### Web Implementation

We have developed a webserver named “CDpred” for the prediction of CD associated peptides. The webserver is implemented by HTML5, JAVA, CSS3 and PHP scripts and compatible on several devices such as iMac, desktop, tablet and mobile. The webserver provides five user-friendly modules such as predict, PQ density, motif scan, protein scan, and design.

## Results

### Positional Conservation Analysis

The specific position of a residue is important for specific role and structure arrangement of a particular peptide or protein. To identify the most significant position of an amino acid residue in the peptide, we perform the positional analysis of CD causing peptides and CD non-causing peptides by using WebLogo (See Figure3). It is worth noting that the first nine locations correspond to peptide N-terminal residues, whereas the latter nine positions correspond to peptide C-terminus. Here, we found that the proline (P) and glutamine (Q) residues are highly prominent at every position while the Phenylalanine (F) and glutamic acid (E) are also found at some positions.

**Figure 3:**
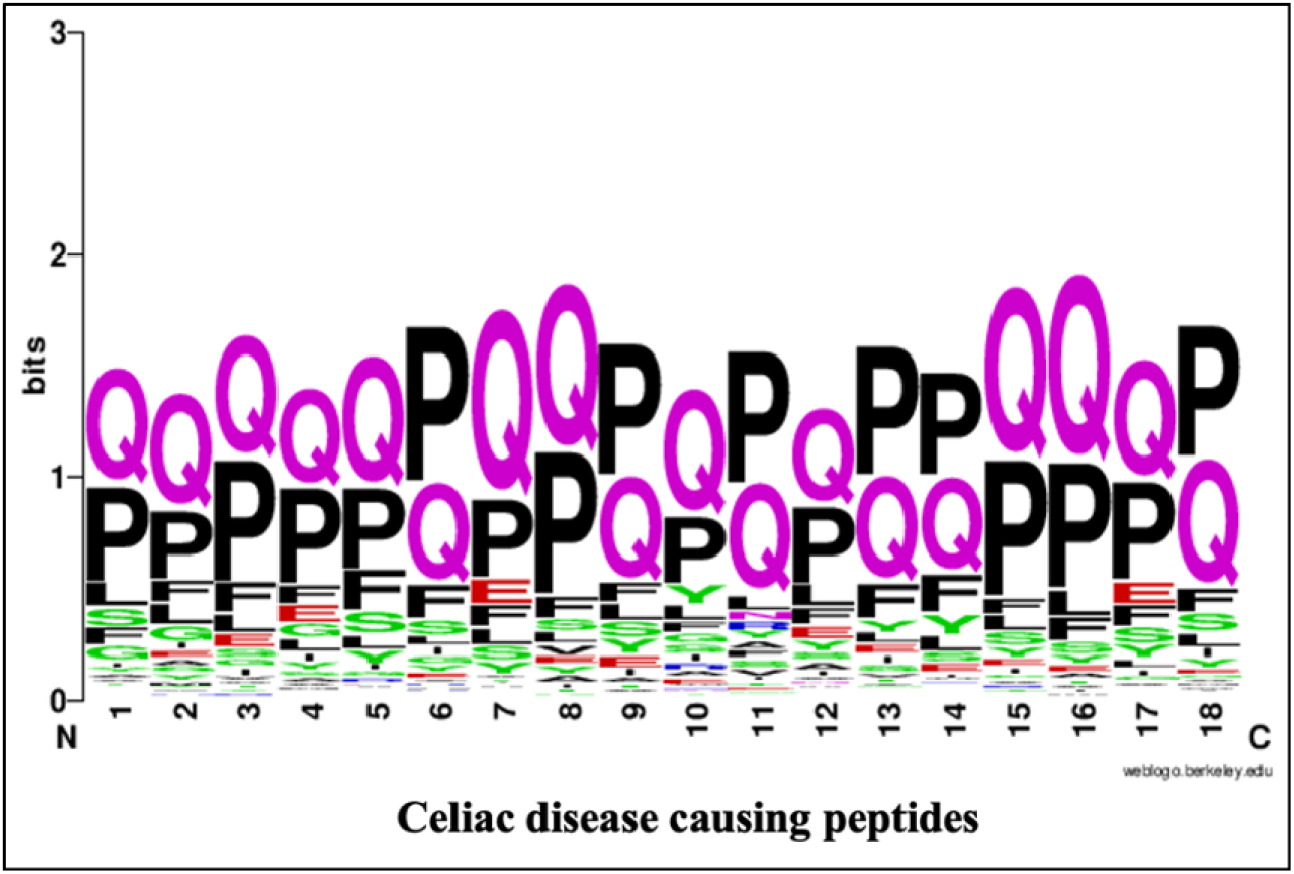
WebLogo of celiac disease-causing peptides.

### Composition Analysis

We compute the amino acid composition for main and alternate datasets. Figure 4 depicts the average composition of CD inducing and non-inducing peptides. In CD causing peptides, the average composition of Proline (P), Glutamine (Q) and Phenylalanine (F) is higher in comparison with disease non-causing peptides, negative random and general proteome.

**Figure 4:**
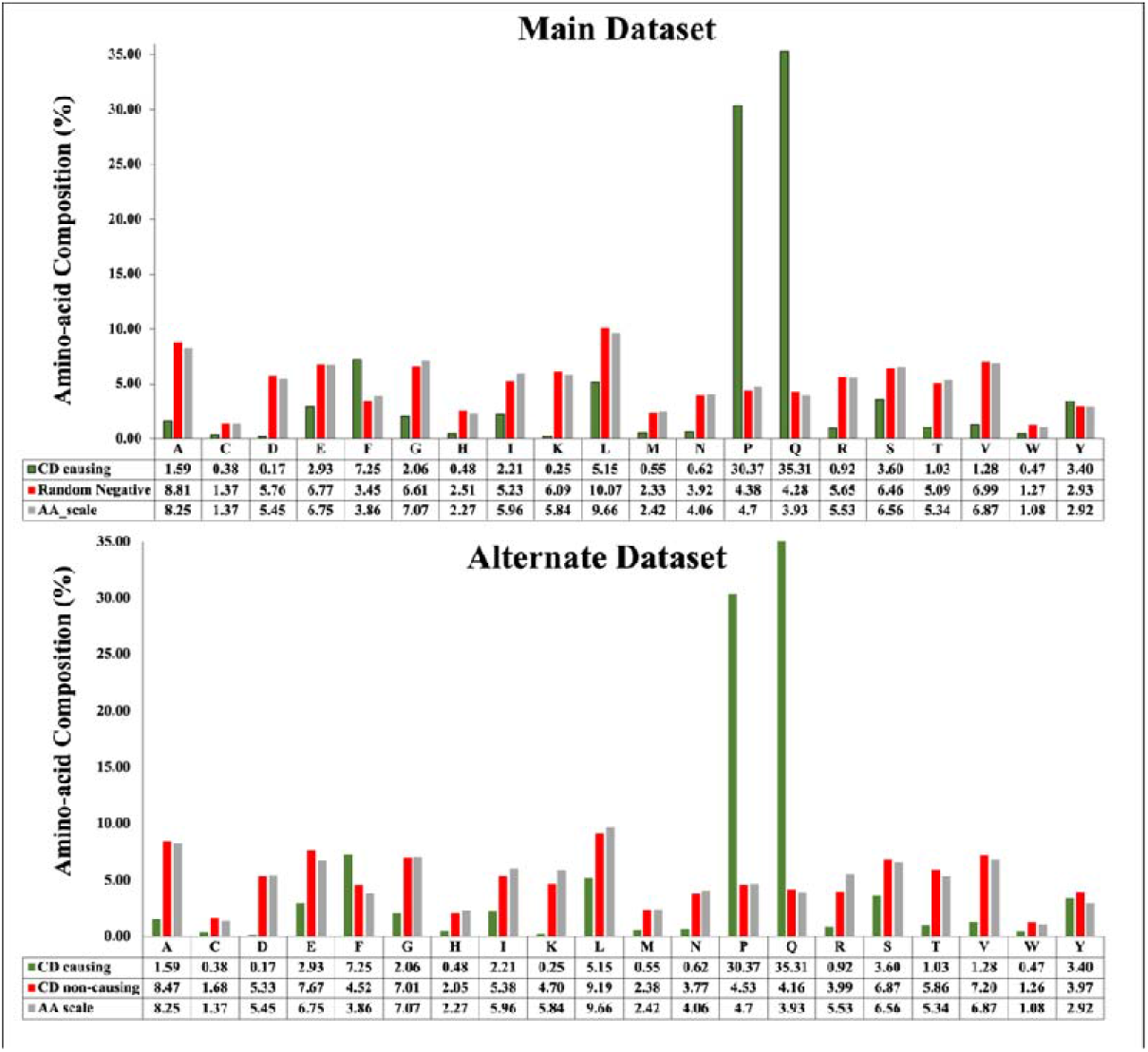
Average amino acid composition of peptides in main dataset and alternate dataset.

### Frequency of HLA alleles

In the past, a number of studies report that celiac disease occurred due to the presence of certain HLA molecules such as HLA-DQ2 and HLA-DQ8 [31]. We have also observed that the maximum gluten immunogenic peptide binders are associated with HLA-DQ2/DQ8 alleles as depicted in Table 1. The complete frequency distribution of HLA-alleles binders of CD causing and non-causing peptides are given in Supplementary Table S1.

**Table 1:**
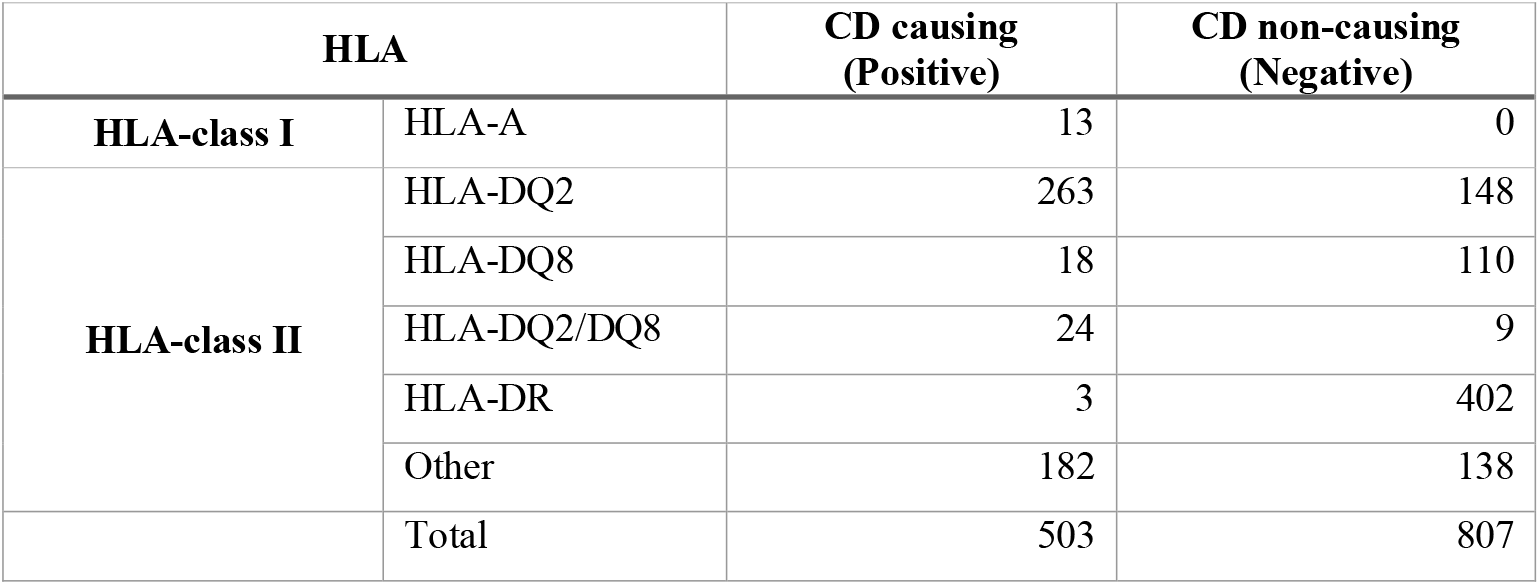
Distribution of HLA alleles in CD causing and non-causing peptides.

### Motif-based Analysis

Motifs are known as the specific regions of a protein sequence which helps to identify the amino acid arrangement shared by a family of protein. The motifs are identified in the CD causing peptide sequences by MERCI program. The MERCI program helps to identify the motif regions in a set of sequences. We utilized the MERCI tool to look for motifs seen only in CD-causing peptides and not in disease non-causing or random peptides. We also looked for motifs found only in disease non-causing and random peptides. Here, we found 50 motifs in CD causing peptides of different length in which P and Q residues are present in abundance in CD causing peptides. We also checked the common motifs found in disease causing, non-causing and random negative peptides. The list of motifs and their occurrence in all the three datasets are given in Table 2.

**Table 2:** Abundance of motifs in CD causing, non-causing and random negative peptides.

| **Motifs** | **CD causing** | **CD non-causing** | **Random Negative** |
| --- | --- | --- | --- |
| **QPF** | 276 | 4 | 0 |
| **QQPF** | 170 | 1 | 0 |
| **PYP** | 120 | 3 | 0 |
| **PEQ** | 56 | 4 | 0 |
| **QPQ** | 350 | 0 | 1 |
| **PQPQ** | 189 | 0 | 1 |
| **QQPQ** | 131 | 0 | 1 |
| **PQL** | 84 | 0 | 1 |

### PQ Density

On performing the compositional and motif analysis, it was found that (P) and (Q) are the most abundant residues in CD-causing peptides as compared to non-causing peptides. In order to classify the peptides based on the PQ density, we have first generated the overlapping patterns of window size ranging from 3 to 9 for each peptide, since 9 was the minimum length of the peptides, and calculated the composition of residues P and Q in each pattern. Each peptide in the dataset is represented by the maximum value of composition for the respective pattern size and found the optimal composition at which we can classify the peptides with balanced sensitivity and specificity. To find the optimal pattern size, we have varied the size from 3 to 9, and found out that window size 5 and 6 performed best among the other sizes for main and alternate datasets, respectively as shown in Table 3.

**Table 3:**
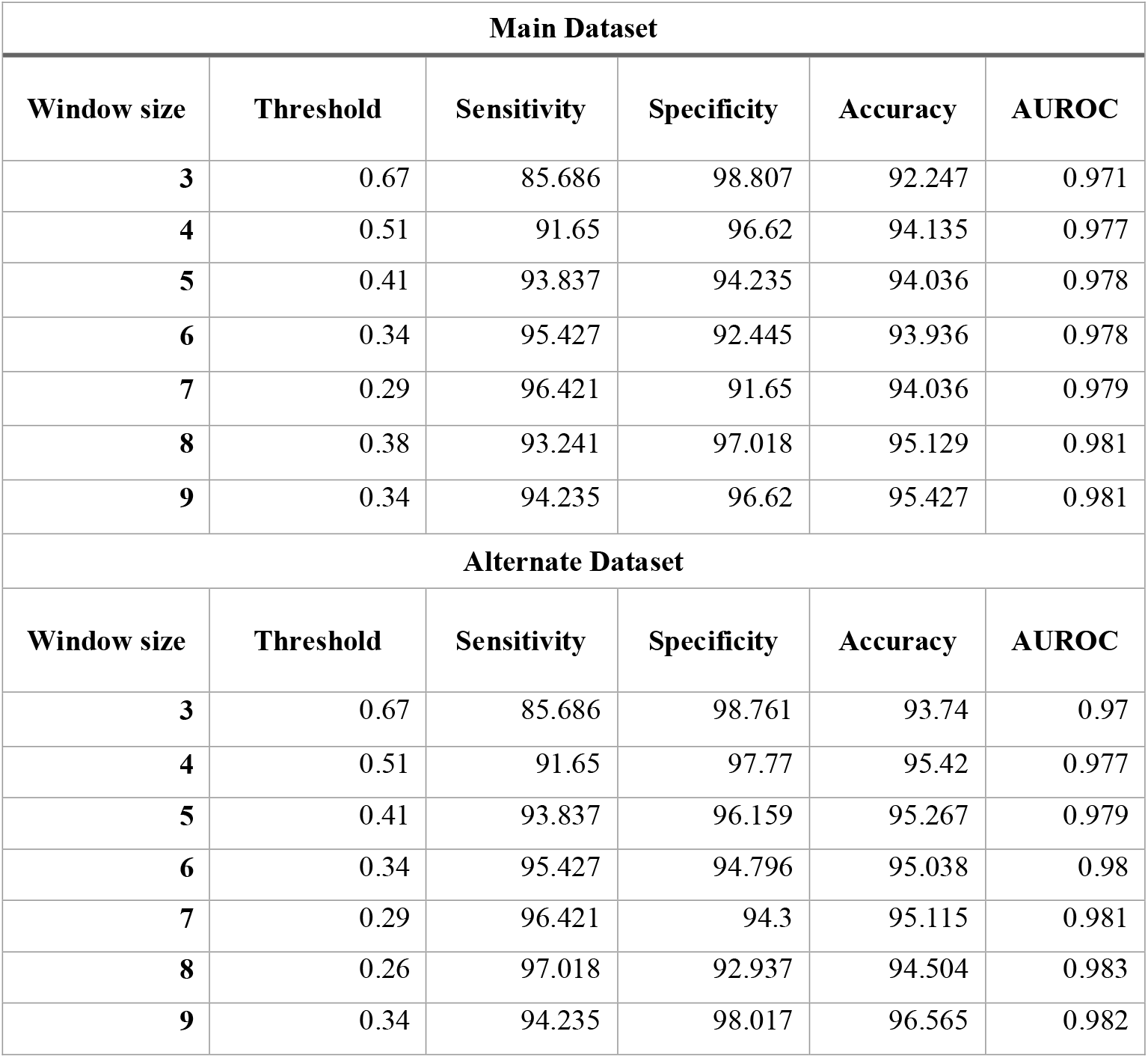
Performance of the PQ Density based method on main and alternate datasets.

### Machine learning based prediction

Various machine learning classifiers such as RF, DT, GNB, XGB, KNN, ETN, SVCN and LR are used to develop a prediction model. For this, we have computed the features of disease causing and disease non-causing peptides using composition-based module of Pfeature.

### Performance of AAC based features

Firstly, we have computed features of amino acid composition, using which we applied different machine learning techniques. As shown in Table 4, ET achieves maximum performance in comparison to other models with AUROC 0.991 and 0.995 and accuracy 96.02 and 97.03 on both training and validation dataset with a good balance of sensitivity and specificity in main data. Similarly, ET achieves maximum performance in comparison to other models with AUROC 0.995 and 0.999 and accuracy 97.519 and 98.092 on both training and validation dataset with a good balance of sensitivity and specificity in alternate data.

**Table 4:** The performance of machine learning classifiers on AAC based features for main and alternate datasets.

|  | Main dataset |  |  |  |  |  |  |  |
| --- | --- | --- | --- | --- | --- | --- | --- | --- |
|  | Training |  |  |  | Validation |  |  |  |
| Classifier | Sens | Spec | Acc | AUROC | Sens | Spec | Acc | AUROC |
| DT | 92.269 | 92.556 | 92.413 | 0.962 | 97.059 | 91 | 94.059 | 0.982 |
| RF | 95.262 | 95.533 | 95.398 | 0.989 | 98.039 | 97 | 97.525 | 0.994 |
| LR | 96.01 | 96.03 | 96.02 | 0.988 | 98.039 | 96 | 97.03 | 0.99 |
| XGB | 95.761 | 95.782 | 95.771 | 0.987 | 98.039 | 93 | 95.545 | 0.995 |
| KNN | 95.262 | 95.285 | 95.274 | 0.986 | 97.059 | 96 | 96.535 | 0.991 |
| GNB | 93.017 | 98.263 | 95.647 | 0.976 | 93.137 | 98 | 95.545 | 0.99 |
| ET | 96.01 | 96.03 | 96.02 | 0.991 | 98.039 | 96 | 97.03 | 0.995 |
| SVC | 95.761 | 95.782 | 95.771 | 0.987 | 97.059 | 96 | 96.535 | 0.991 |
|  | Alternate dataset |  |  |  |  |  |  |  |
|  | Training |  |  |  | Validation |  |  |  |
| Classifier | Sens | Spec | Acc | AUROC | Sens | Spec | Acc | AUROC |
| DT | 92.537 | 92.57 | 92.557 | 0.968 | 94.059 | 99.379 | 97.328 | 0.99 |
| RF | 97.015 | 97.368 | 97.233 | 0.995 | 98.02 | 97.516 | 97.71 | 0.998 |
| LR | 96.269 | 96.285 | 96.279 | 0.99 | 97.03 | 96.273 | 96.565 | 0.987 |
| XGB | 97.015 | 97.059 | 97.042 | 0.992 | 99.01 | 93.168 | 95.42 | 0.998 |
| KNN | 95.771 | 95.975 | 95.897 | 0.992 | 98.02 | 95.652 | 96.565 | 0.995 |
| GNB | 92.537 | 97.059 | 95.324 | 0.977 | 96.04 | 96.273 | 96.183 | 0.983 |
| ET | 97.512 | 97.523 | 97.519 | 0.995 | 98.02 | 98.137 | 98.092 | 0.999 |
| SVC | 97.015 | 96.904 | 96.947 | 0.993 | 98.02 | 96.894 | 97.328 | 0.996 |

**Table 5:**
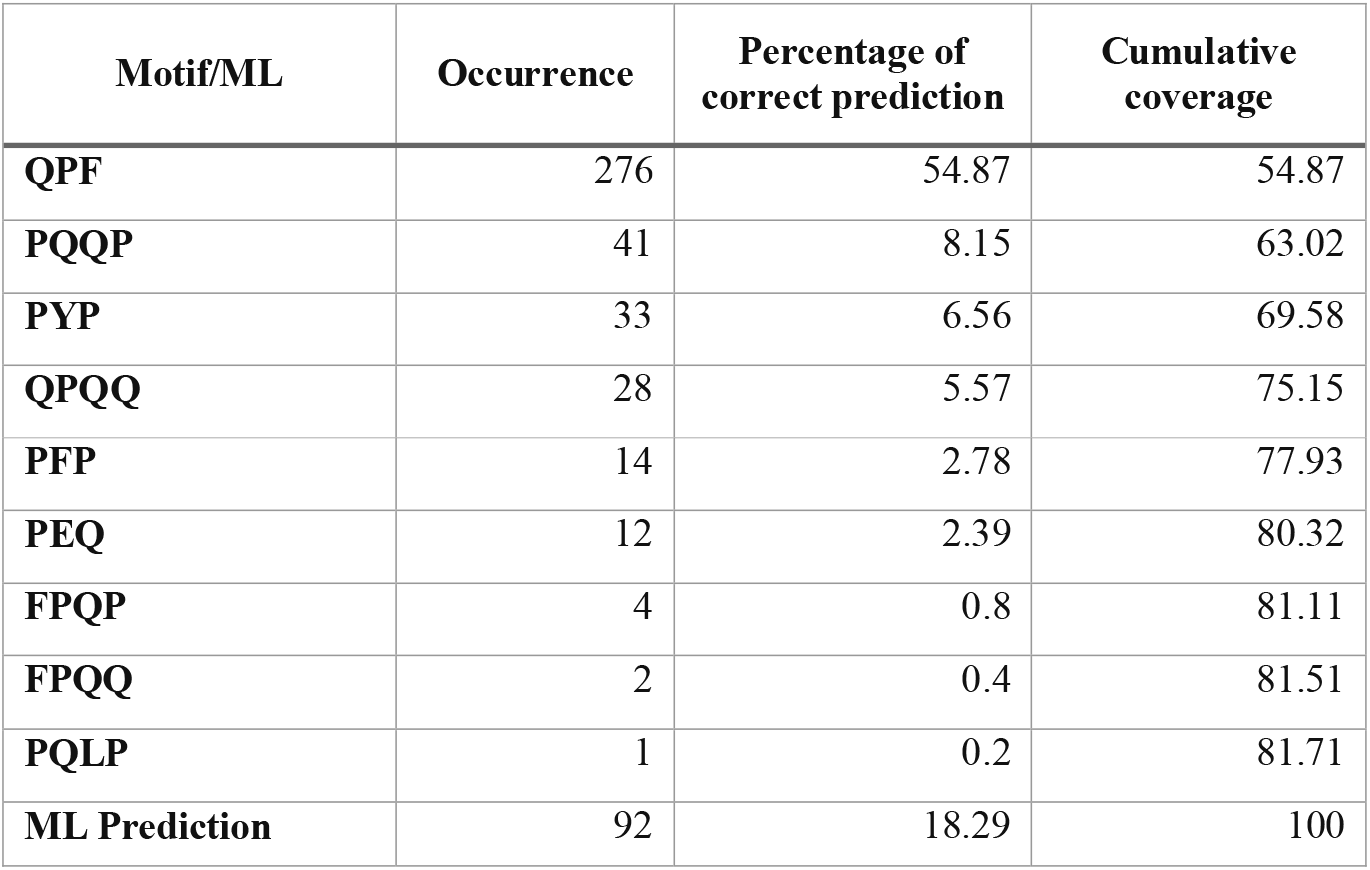
The table shows the occurrence of motif in positive sequences with percentage of correct prediction and cumulative coverage.

### Performance of Ensemble model

In ensemble method, first we used the motif-based approach by identifying the coverage of motifs in the given protein/peptide sequences. Our motif-based approach achieves 81.71% accuracy in the independent dataset. The rest sequences, which were not predicted using motif-based approach, were covered by using the machine learning method. By combining both approaches we achieve the highest performance of 100% accuracy on independent dataset. Our ensemble method is the best approach for predicting the CD associated peptides.

### Services to Scientific Community

We design a user-friendly prediction web server that incorporates several modules to determine CD-causing peptides in order to serve the scientific community. The prediction models which used in the study are implemented in the web server. Based on the prediction models’ score at a different threshold, users can predict whether a query peptide causes CD or not. The web server comprises five major modules 1) Prediction, 2) PQ Density, 3), Motif 4) Scan and 5) Design. The user can classify CD-causing peptides from disease non-causing peptides using the ‘Predict’ module. The “PQ Density” module used to calculate PQ content in a given query sequence based upon the window size. Users can map or scan CD-causing motifs in the query sequence using the “Motif” module. We used the MERCI software to extract themes from CD-causing peptides that had been empirically confirmed. The “Scan” module was used to scan the amino-acid sequence for CD-causing areas. The user can generate all potential analogs of the input sequence using the “Design” module. The positive and negative datasets utilized in this work are also available for download, and the peptide sequence are available in FASTA format. HTML, JAVA, and PHP scripts were used to create the web server CDpred https://webs.iiitd.edu.in/raghava/cdpred/. The server is user-friendly and compatible with a variety of devices, including computers, Android phones, iPhones, and iPads. In addition, we provided a standalone package in the form of a Docker container.

## Discussion & Conclusion

Celiac disease is a gluten-sensitive enteropathy caused due to the chronic, genetically predisposed, and autoimmune condition with a wide spectrum of clinical manifestations brought on by consuming gluten [32]. CD mainly effect the immune system of the patients and causes a number of diseases such as cirrhosis, autoimmune hepatitis, diabetes mellitus, gluten ataxia, peripheral neuropathies, etc [33,34]. During disease condition, enormous amount of CD4+ T cell response act against gluten peptides which are presented by HLA-DQ molecules [35]. In such situation, our immune system secretes excess amount of cytokines and gluten protein specific antibodies. The only effective life-long treatment of this disease is gluten-free diet. Due to increased number of cases in worldwide a number of gluten-free products are available for celiac susceptible people [17,36]. Thus, it is essential to identify or eliminate gluten immunogenic peptides from the food products which can induce the celiac disease and sensitive to celiac patients.

In this study, we have made a systematic attempt for the prediction of peptides responsible for causing the disease. We have collected the dataset from IEDB and Swiss-Prot databases. We have created two datasets for the analysis and prediction of CD causing peptides. Here, we observed that residues (P and Q) are highly abundant in CD causing peptides in comparison with negative and random peptides. The similar findings are supported by the previous studies where they have shown the abundance of P and Q amino acids in gluten proteins [37,38]. From the motif-based approach we identified motifs [QPQ, QPF, PQPQ, QQPF, QQPQ, PYP], which are highly conserved in CD causing peptides in comparison with negative dataset. So, we performed PQ based analysis where we calculate the abundance of PQ residues in the CD causing and non-causing peptides.

In addition, we have developed prediction models using amino-acid composition-based features. We achieved maximum performance with AUROC of 0.99 on the training and validation datasets, respectively. We have also developed an ensemble method by combining both motif-based approach and machine learning based models. This ensemble approach provides the 100% accuracy on independent dataset. In addition, we have developed a webserver named CDpred (https://webs.iiitd.edu.in/raghava/cdpred/), standalone package (https://webs.iiitd.edu.in/raghava/cdpred/standalone.php) and GitLab (https://gitlab.com/raghavalab/cdpred) for the prediction of CD causing peptides.

## Authors’ contributions

RT, AD and GPSR collected and processed the datasets. RT, SP and GPSR implemented the algorithms and developed the prediction models. RT, AD, SP and GPSR analysed the results. RT and SP created the back-end of the web server the front-end user interface. RT, AD, and GPSR penned the manuscript. GPSR conceived and coordinated the project. All authors have read and approved the final manuscript.

## Acknowledgements

Authors are thankful to the Department of Bio-Technology (DBT) and Department of Science and Technology (DST-INSPIRE) for fellowships and the financial support and Department of Computational Biology, IIITD New Delhi for infrastructure and facilities.

## Abbreviations

CD: Celiac Disease
HLA: Human leukocyte antigens
CXCR3: Chemokine receptor 3
tTG: Tissue transglutaminase
sIgA: Secretory Immunoglobulin A
IEDB: Immune Epitope Database
AUROC: Area under receiver operator curve
DT: Decision Tree
RF: Random Forest
SVC: Support Vector Classifier
XGB: XGBoost
LR: Logistic Regression
ET: Extra Tree classifier
KNN: k-Nearest Neighbors
GNB: Gaussian Naive Bayes

